# Microglia extract neuronal proteolytic organelles via skoupocytosis

**DOI:** 10.64898/2026.07.22.740182

**Authors:** Renee Pepper, Jacqueline M. Griswold, Frances Middleton-Davis, Sarah Syed, Yuuta Imoto, Lani Tran, Eunbin Park, Yu Kang T. Xu, Dwight Bergles, JoAnn Buchanan, Shigeki Watanabe

**Affiliations:** Department of Cell Biology, Johns Hopkins School of Medicine, Baltimore, MD, USA; Department of Developmental Neurobiology, Division of Neuronal Cell Biology, St Jude Children’s Research Hospital, Memphis, TN, USA; Solomon H. Snyder Department of Neuroscience Johns Hopkins School of Medicine, Baltimore, MD, USA; Department of Neurobiology, Howard Hughes Medical Institute, Harvard Medical School, Boston, MA, USA; Kavli Neuroscience Discovery Institute, Johns Hopkins University, Baltimore, MD, USA; Allen Institute for Brain Science, Seattle, WA, USA; Department of Neurology and Neurological Sciences, Stanford University, Stanford, CA, USA

## Abstract

Neurons face unique challenges in maintaining protein homeostasis due to their tortuous morphology and extended processes. Proteolytic organelles are typically transported retrogradely towards their soma for degradation where lysosomes are enriched, but some organelles are larger than these neuronal processes, questioning how these organelles are degraded. Here we show that microglia, the resident immune cells of the brain, extract proteolytic organelles from neurons both *in vitro* and *in vivo*. Microglia make transient contact with neuronal membranes where proteolytic organelles are stationed beneath. At these sites of interaction, microglia pinch off a small portion of the neuronal process containing the organelle, leaving the rest of the process intact. We term this process ‘skoupocytosis’ after the Greek word for garbage. Phosphatidylserine (PS) lipase ABHD16a accumulates near these proteolytic organelles and converts PS into lyso-PS to initiate microglial skoupocytosis. Skoupocytosis bypasses the need for retrograde organelle transport, providing homeostatic advantages for neurons that must maintain function in processes that extend extraordinarily long distances from the cell soma.

## Main

Neurons face a unique problem maintaining protein homeostasis due to their extremely polarized morphology and high rates of membrane and protein turnover during synaptic communication. Neurons are also post-mitotic and cannot dilute their proteolytic burden with division, so interruptions to neuronal proteostasis have disastrous effects on neuronal health^1–3^. Thus, neurons are equipped with several proteolytic pathways. Misfolded or damaged proteins, and organelles destined for degradation are enveloped by double membrane phagophore to generate autophagosomes in a process known as autophagy^4–6^. Transmembrane proteins undergo endosomal sorting mediated by hepatocyte growth-factor tyrosine kinase substrate (HRS) following endocytosis and are sorted into multi-vesicular bodies (MVBs)^7,8^. These highly active pathways maintain homeostasis of neuronal proteins to sustain signaling and enable plasticity.

Proteolytic organelles are thought to be transported back to the cell soma for degradation. Seminal studies performed in cultured neurons demonstrated that autophagosomes and MVBs are generated in distal axons and mature into autolysosomes and amphisomes during retrograde transport^9–12^. Eventually, they fuse with lysosomes, which are enriched in the proximal axon and the cell soma^9–12^. However, proteolytic organelles can be larger than axons, particularly in unmyelinated axons that exhibit pearls-on-a-string morphology where the diameter of the string region can be as small as ∼60 nm^13^, raising the question of whether all proteolytic organelles follow this canonical pathway.

Here we describe a new pathway by which microglia contribute to neuronal proteostasis. Specifically, we found larger HRS-positive proteolytic organelles such as amphisomes, MVBs and autophagosomes from neurons are indeed stationary when neurons are cultured alone. However, upon addition of microglia into the culture, microglia take up these larger proteolytic organelles from neurons. *In vivo* 2-photon imaging revealed that this same process also occurs in pyramidal neurons of the somatosensory cortex. In electron micrographs, microglia appear to engulf a portion of synapses that contain proteolytic organelles, suggesting remarkable spatial specificity. We further demonstrate that a/ß hydrolase domain-containing (ABHD16a), a phosphatidylserine (PS) lipase, is enriched at sites containing proteolytic organelles, along with externalized PS, a known ‘eat me’ signal that triggers microglial engulfment, and that lyso-PS generated by ABHD16a is necessary for skoupocytosis. Together, these results reveal that microglia assist in local removal of proteolytic organelles from neuronal processes of which we termed Skoupocytosis after the Greek word for garbage ‘Skoupidia’.

## Results

### Large proteolytic organelles in neurons are stationary

The trafficking of proteolytic organelles is a dynamic process influenced by neuronal activity, stress, age and disease^14,15^. The speed and direction of autophagic organelles is influenced by different autophagy markers^16,17^, however the effect of organelle size on their dynamics and fate is less characterized. To address this question, we tracked proteolytic organelles in axons of hippocampal neurons removed from embryonic day 18 (E18) C57BL/6N mice and maintained in culture. To visualize proteolytic organelles, HRS-HaloTag (Halo) was expressed using lentivirus on days *in vitro* (DIV) 9. HRS initiates endosomal sorting, resulting in the labelling of proteolytic organelles. On DIV14, these neurons were labelled with the Janelia dye JFX-646 for 30 min, and live-cell imaging was performed at 200 ms intervals over 100 s (Fig. 1a,b). We quantified the net displacement (µm) of HRS-Halo+ organelles categorized by size (µm^2^). We observed that small HRS+ organelles ranging from 0.1-0.2 µm^2^ were most abundant and mobile, undergoing a net displacement of 0.39 µm over 100 s. Organelles ranging from 0.2-0.4 µm^2^ moved the furthest averaging a net displacement of 0.46 µm (Fig. 1c). As the average size of the organelle increased, the net displacement decreased, with organelles with an area greater than 1.6 µm^2^ moving significantly less than those between 0.2-0.4 µm^2^ (Ordinary one-way ANOVA with Tukey’s multiple comparisons, p = 0.0069, n = 3). Of particular interest, some large organelles remained stationary in regions of the axon where smaller organelles were translocated (Fig. 1a,b), suggesting that the size of oragnelles may contribute to their ability to traffic along neurites.

**Fig. 1:**
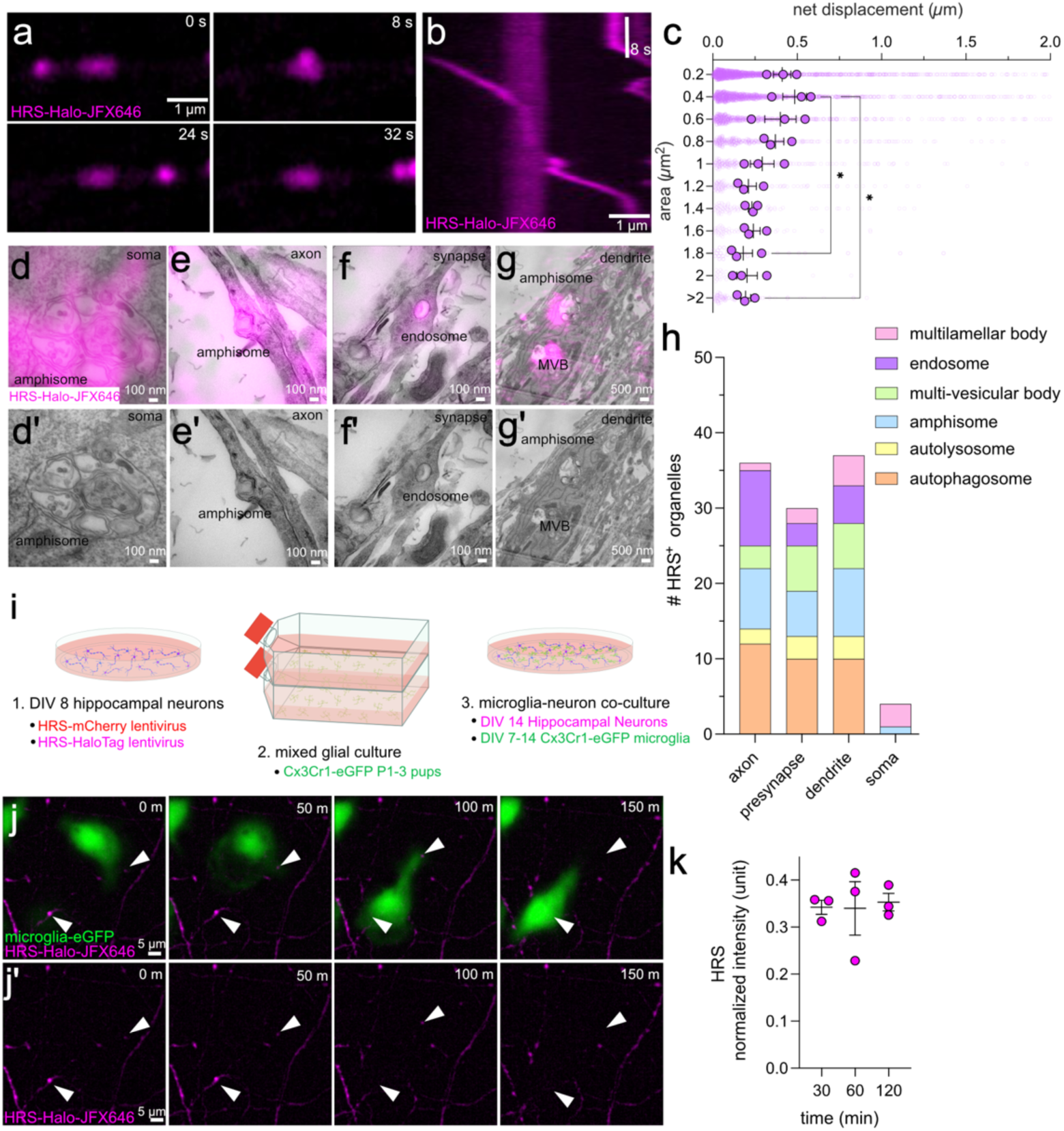
Large, stationary proteolytic organelles in neurons are taken up by microglia *in vitro*. (a) Time-lapse images of HRS-Halo-JFX646-labeled organelles in DIV21 hippocampal neurons 5fps (200 ms intervals). Scale bar 1 µm. (b) Kymograph from time-lapse imaging of HRS-Halo-JFX646-labeled organelles in DIV21 hippocampal neurons. Scale bar 1 µm and 8 seconds. (c) Net displacement (µm) of HRS-Halo-JFX646 organelles by area (µm²). n = 3 coverslips per N = 3 independent hippocampal cultures. Statistical analysis using Ordinary one-way ANOVA with Tukey’s multiple comparisons test indicates significant differences between the groups as indicated by the asterisk (* p<0.05), with data represented as mean ± SEM. (d-g) Electron micrographs overlayed with correlative fluorescent signal from HRS-Halo-JFX646-labeled organelles from different cellular compartments featuring: (d) amphisome at soma, (d) amphisome in axon, (e) endosome at presynapse, (f) amphisome and multi-vesicular body in dendrite. Scale bars are 100 µm for (c–e) and 500 µm for (f). (h) Bar graph quantifying the number of HRS^+^ organelles present in subcellular compartments of the neuron (axon, presynapse, dendrite, soma) categorized by type: multilamellar body (red), endosome (orange), multi-vesicular body (yellow), amphisome (cyan), autolysosome (purple), and autophagosome (green). n = 3 independent hippocampal cultures. (i) Experimental design schematic: (1) DIV 8 hippocampal neurons infected with HRS-mCherry lentivirus and HRS-Halo lentivirus; (2) mixed glial culture from Cx3Cr1-eGFP P1-3 pups; (3) co-culture of DIV 14 hippocampal neurons with DIV 7-14 Cx3Cr1-eGFP microglia. (j) Time-lapse images of microglia (green, Cx3Cr1-eGFP) engulfing HRS-Halo-JFX646-labeled organelles (magenta) from DIV14 hippocampal neurons (1-minute imaging intervals). j’) same images with no microglia shown. White arrows indicate HRS-Halo-JFX646-labeled organelles that are engulfed. Scale bar 5 µm. (k) Normalized intensity of HRS-mCherry (arbitrary units, normalized to min/max) internalized by microglia at 30, 60, 120 mins. Data shown as mean ± SEM. n = 3 independent hippocampal cultures.

To determine the identity of the large stationary organelles, we performed correlative light and electron micrscopy (CLEM) of HRS-Halo+ puncta, as described previously^16^. Besides endsomes (Fig. 1f), autophagosomes, autolysosomes and multi-vesiclular bodies (MVBs) (Fig. 1g), as previously reported, HRS were also localized to amphisomes (Fig. 1d,e,g) and multilamellar bodies (Extended Data Fig. 1)^8,19,20^. The prevalence of these proteolytic organelles was largely similar across different cellular compartments including axons, dendrites and presynapses, but only amphisomes and multilamellar bodies were observed at the cell soma (Fig. 1h, Extended Data Fig. 1). These data suggest that large stationary HRS-Halo+ puncta are proteolytic organelles.

### Microglia take up proteolytic organelles from neurons *in vitro*

The presence of large stationary proteolytic organelles in axons and dendrites raises new questions about how neurons prevent their expansion and accumulation, which could eventually impair the ability of the cell to clear misfolded and modified proteins. While some lysosomes are present in axons and dendrites, the majority reside in the soma^9,10,21^. Historically, the majority of studies exploring proteolytic organelle trafficking in neurons have used *in vitro* culture systems where neurons are cultured alone or with sparse astocytes, but not with professional phagocytic cells of the mammalian central nervous system (CNS) like microglia. Thus, we isolated microglia from mixed glial cultures from *Cx3Cr1-eGFP* P0-P3 mouse pups (B6.129P2(Cg)-*Cx3cr1^tm1Litt^*/J) using the shake-off method^22^, and seeded them with DIV14 hippocampal neurons expressing HRS-Halo labelled with JFX-646 dye 30 min prior to imaging (Fig. 1i). To our surprise, we observed microglia extending processes to regions of the neurons where HRS+ puncta were located and eventual transfer of these puncta into the cell (Fig. 1j,k, Supplementary Video 1), suggesting that microglia may be able to recognize the presence of proteolytic organelles in neurons and extract them. To confirm their internalization, we co-cultured neurons expressing HRS-mCherry (more resistant to low pH and degradation) and microglia from wild-type mice and performed immunocytochemistry. At all three timepoints HRS-mCherry puncta (visualized with a anti-dsRed antibody), were observed within microglia (Fig. 1k). Together, these data suggest proteolytic organelles can be removed from neurons by microglia *in vitro*.

### Microglia take up proteolytic organelles from neurons *in vivo*

The transcriptomic and molecular features of microglia are particularly sensitive to their environment, and culturing microglia has shown to induce considerable changes to their cellular properties^23^. To test whether proteolytic organelles are taken up by microglia under more physiological conditions, we performed *in vivo* 2-photon imaging (Fig. 2). To express fluorescently tagged HRS *in vivo* we generated an adeno-associated virus (AAV) using a PHP.eB capsid with a human Synapsin promotor to express HRS-mCherry specifically in neurons (Fig. 2a)^24,25^. This AAV was injected intracerebroventricularly into *Cx3Cr1-eGFP* mice (tg/+) at postnatal day 0-2 (P0-2). Upon reaching adulthood (postnatal days 60-120, P60-120) mice received a 3 mm diameter craniotomy to implant a cranial window over their primary somatosensory cortex (see Methods).

**Fig. 2:**
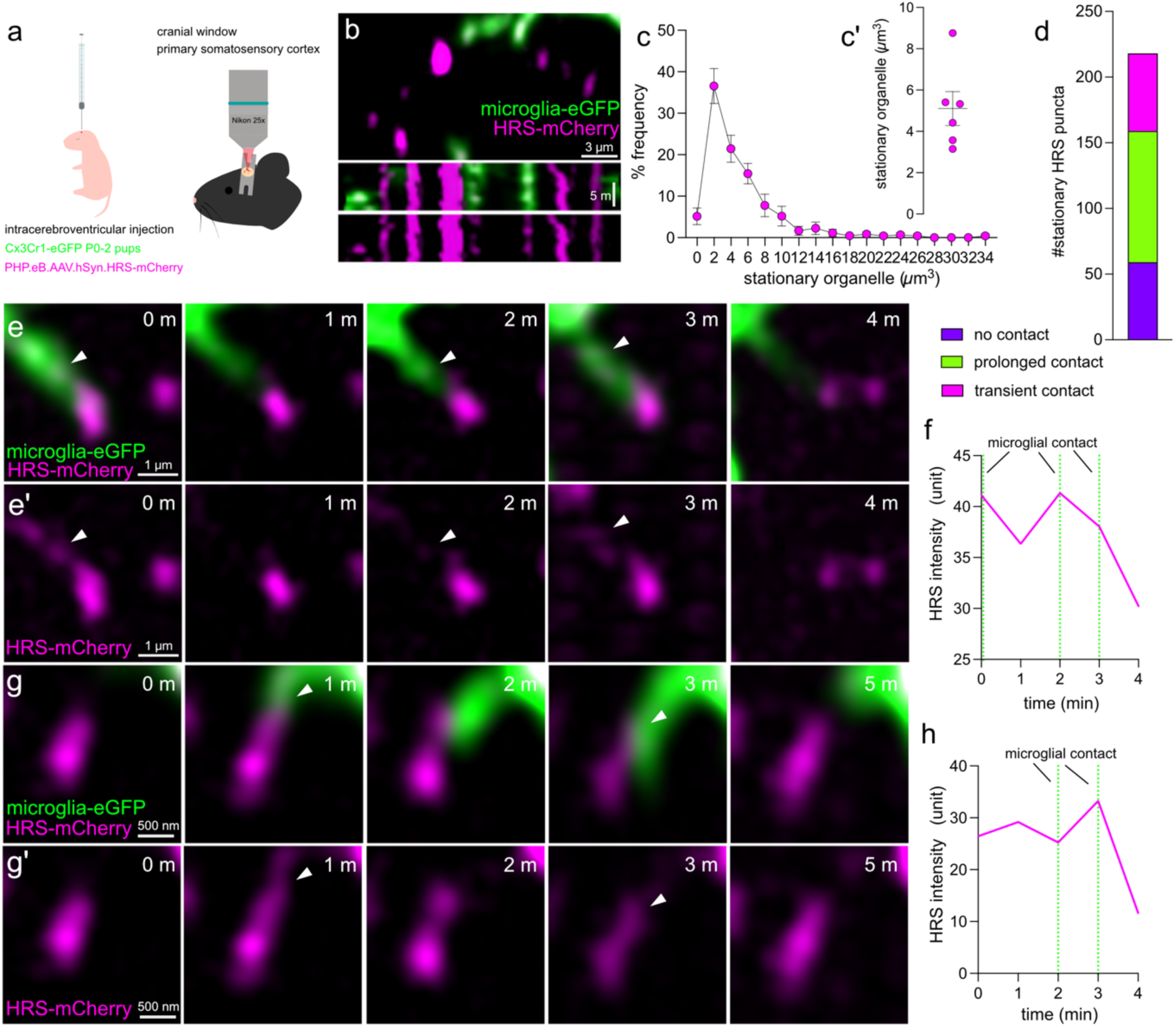
Microglia take up proteolytic organelles from neurons *in vivo*. (a) Experimental design schematic: (1) Cx3Cr1-eGFP P0-2 pups injected with PHP.eB.AAV.hSyn.HRS-mCherry unilaterally into intracerebroventricular cavity; (2) Implantation of cranial window over primary somatosensory cortex for 2-photon *in vivo* imaging 100-150 µm below coverslip in layer II/III. (b) Average intensity projection of time-lapse images from Cx3Cr1-eGFP (green) and neuronal HRS-mCherry^+^ (magenta) organelles. Kymograph from time-lapse imaging of HRS-mCherry^+^ labeled organelles. Scale bar 3 µm and 5 minutes. (c) Histogram representing mean percentage frequency of stationary HRS-mCherry^+^ organelle volume (µm^3^). (c’) Mean stationary HRS-mCherry^+^ organelle volume (µm^3^) per animal. Data shown as mean ± SEM. n=6 mice. (d) Number of HRS-mCherry^+^ stationary organelles either (i) contacted (purple), (ii) prolonged contact (green), (iii) transient contact (magenta). n=6 mice. (e) Time-lapse images of microglia from Cx3Cr1-eGFP (green) taking up neuronal HRS-mCherry^+^ (magenta) organelles. 1-minute imaging intervals. Scale bars represent 1 µm. Representative images from P70 mouse. (e’) same images without microglia shown. (f) Hrs-mCherry-labeled organelle intensity (arbitrary units, normalized to min/max) over time (magenta) during microglial contact. Dotted green line indicates microglial contacts. (g) Time-lapse images of interactions between microglia from Cx3Cr1-eGFP (green) mouse with neuronal HRS-mCherry (magenta) labeled organelles. 1-minute imaging intervals. Scale bars represent 500 nm. Representative images from P70 mouse. (g’) same images without microglia shown. (h) HRS-mCherry-labeled organelle intensity (arbitrary units, normalized to min/max) over time (magenta) during microglial contact. Dotted green line indicates microglial contacts.

Volumetric imaging in layer II/III of these young mice revealed the presence of HRS-mCherry+ structures contained within the cell body and processes of microglia (Supplementary Video 2a). There were also two categories of HRS puncta that were not contained within microglia, including small fast-moving and/or transient HRS puncta most of which were below 1 µm^3^ (Supplementary Video 2a-b). Due to constraints in optical and temporal resolution it was not possible to accurately quantify the dynamics of these small organelles. There were also large stationary puncta similar to those observed in culture that were stationary over the course of the 15-minute imaging periods (Fig. 2b; Supplementary Video 2b). The volume of stationary organelles ranged from around 1-34 µm^3^ with an average size 5.1 µm^3^ (Fig. 2c, d). Stationary organelles were then manually scored for whether they were contacted by microglia, how many contacts they received and whether transfer of HRS signal to microglia occurred. On average, 77% of stationary HRS puncta were contacted by microglia during the 15-minute imaging period, and of these 54% associated for the duration of the imaging period (Fig. 2d). Contact between microglia and stationery HRS puncta occurred in two ways. Some microglia processes were closely associated alongside HRS puncta for the entire imaging period, and HRS intensity could be seen increasing inside the adjacent microglia over time (Supplementary Video 2e). In the second mode of interaction, microglia extended processes towards stationary HRS puncta, transiently contacted these puncta before the HRS signal was translocated from neurites to the microglial cell (Fig. 2e, g; Supplementary Video 2c-d). Consequently, the HRS-mCherry intensity in neurons was reduced following microglial contact (Fig. 2f, h), suggesting removal of the organelle. Remarkably, these interactions were rarely associated with uptake of entire HRS signals. Instead, only some of the HRS signal was transferred at microglia contacts with stationary organelles, leaving the remaining HRS signal in the neurons (Fig. 2e, g). Together, these data suggest that microglia specifically reach out to parts of neurites where proteolytic organelles accumulate and assist in their removal.

### HRS uptake is independent of synaptic pruning

Although 2-photon *in vivo* imaging provided evidence for the transfer of proteolytic organelles from neurons to microglia, the prevalence and frequency of such events are difficult to quantify due to the insufficient spatial and temporal resolutions of this imaging modality. Furthermore, microglia are known to engulf entire synapses to refine circuits^26–30^, and thus, HRS-mCherry may be removed coincidentally during synaptic pruning. To quantify the number of contacts between microglia and proteolytic organelles or synapses, superresolution imaging (SoRa) was performed in *Cx3Cr1-eGFP* mice expressing PHP.eB.AAV.hSyn.HRS-mCherry as described above after perfusion fixation at P16, P60 and P240 (Fig. 3). Presynapses were marked with Bassoon antibody and quantified the fluorescence signals in the hippocampus (HC) and in the somatosensory cortex (SS) (Fig. 3a-c). The average volume of HRS puncta per image remained relatively constant across these three time points (Fig. 3d). Some HRS puncta were found next to the Bassoon puncta, but these signals did not fully overlap (Fig. 3c). Approximately 13% of synapses contained HRS puncta at P16 and P60 (Fig. 3e). The prevalence and intensity of these HRS puncta at synapses increased slightly at P240 (Fig. 3e, f), indicating potential higher proteolytic activity in older mice.

**Fig. 3:**
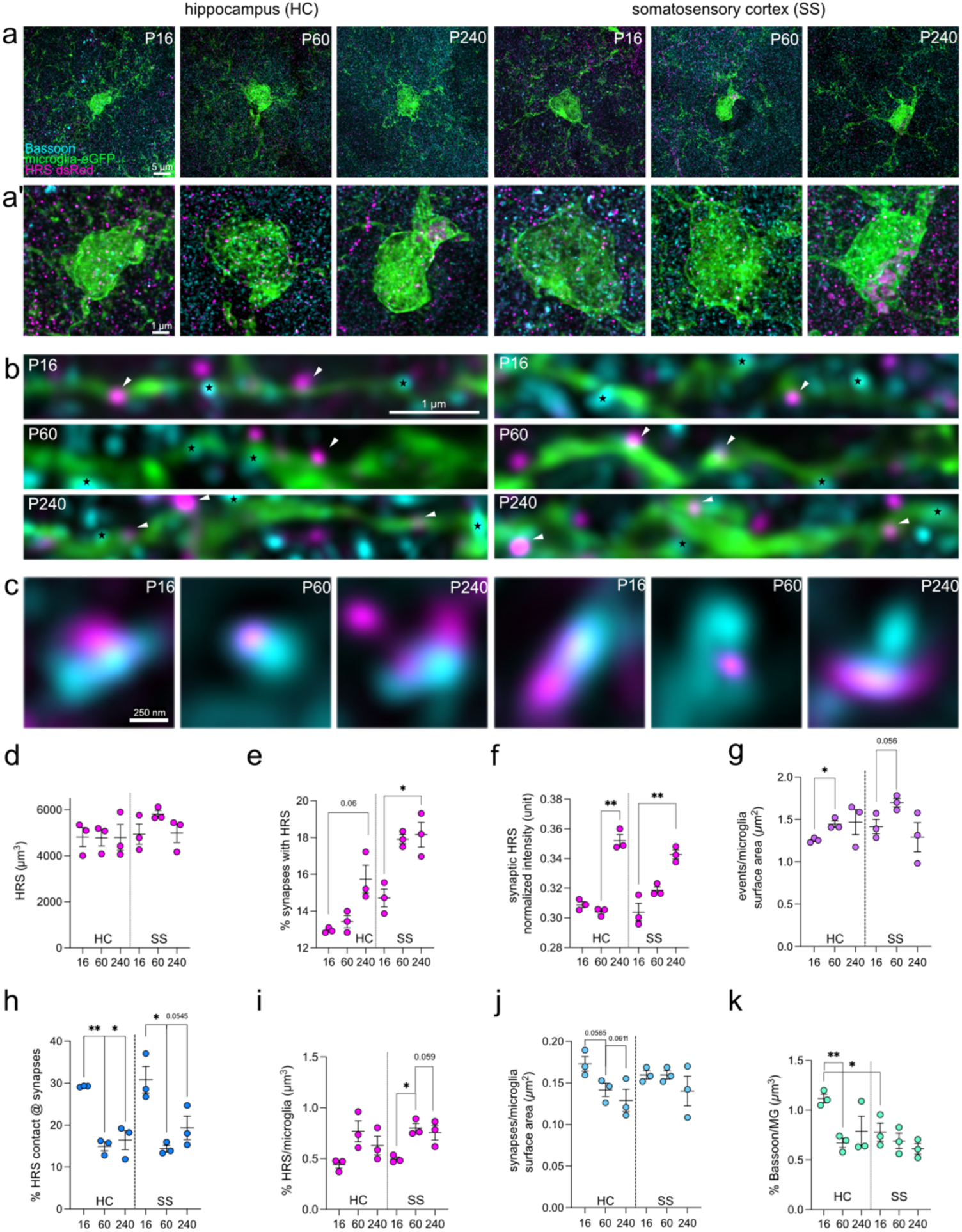
Proteolytic organelle density and uptake vary based on region and age. (a) Representative immunofluorescence images take from Cx3Cr1-eGFP (green) mice expressing neuronal HRS-mCherry (dsRed antibody to HRS-mCherry, magenta) and presynaptic active zones (Bassoon antibody, cyan) at fixed time points of P16, P60, and P240 in the hippocampus (HC) and somatosensory cortex (SS). (a’) zoomed in images of (a). (b) Images show microglia (green) contacting HRS (magenta) indicated by white arrows, and presynaptic active zone (Bassoon antibody, cyan) indicated by black star. Scale bars represent 1 µm. (c) Images show HRS (magenta) near active zone marked by Bassoon antibody (cyan). Scale bar 250 nm. (d) Quantification of HRS (µm^3^) density. (e) Percentage of presynapses associated with HRS. (f) Normalized HRS intensity associated with synapses (Bassoon^+^) (arbitrary units, normalized to min/max) (g) Number of contacts with HRS per 10 µm^2^ microglia surface area. (h) Percentage of the contact in (g) that are near the active zone. (i) Percentage HRS volume (µm^3^) of microglia volume (µm^3^). (j) Number of contacts with Bassoon per 10 µm^2^ microglia surface area. (k) Percentage Bassoon volume (µm^3^) of microglia volume (µm^3^). n=7-15 microglia from n=3 mice. Each dot represents 1 animal. Data represented as mean ± SEM. Statistical analysis using Welch’s t-test, p-value represented in figure. * p < 0.05, ** p < 0.001.

Next, we quantified microglia interactions with 1) HRS puncta, 2) Bassoon puncta, and 3) HRS puncta near Bassoon puncta. Interactions were considered as ‘contact’ when the overlap of the signals was greater than 1 pixel and ‘internalized’ if the HRS or Basson puncta were 100% contained within GFP signals. Contact number was normalized to microglia surface area (µm^2^) per image. The number of HRS puncta contacted by microglia was approximately 1.2 per 10 µm^2^ surface area of microglia at P16 and increased to 1.4 at P60 and remained elevated at 1.4 at P240 (Fig. 3b, g) (unpaired t-test with Welch’s correction, p = 0.0218, n = 3). Of these contacts, 29% were found near the presynaptic active zone (Fig. 3h), suggesting that this process could occur around presynapses. The volume of HRS puncta internalized by microglia increased at P60 and remained elevated at P240 (Fig. 3i). By contrast, the number of microglia contact with Basson puncta was higher (approximately 1.9 events per 10 µm^2^ surface area of microglia) at P16 and decreased to 1.4 at P60 and further down to 1.2 at P240 (Fig. 3j). Likewise, the amount of Bassoon puncta internalized by microglia was highest at P16 and decreased at later time points (Fig. 3k) (unpaired t-test with Welch’s correction, p = 0.0024, n = 3), potentially representing synaptic pruning, given its higher prevalence earlier in life^26–30^. Similar trends were observed in the somatosensory cortex (SS) (Fig. 3). Together, these results suggest that microglia can detect areas along neurites where proteolytic organelles are located and potentially internalize HRS-positive organelles from neurons independent of their role in synaptic pruning.

### Microglia nibble the areas that contain proteolytic organelles – skoupocytosis

Superresolution imaging provided the spatial specificity of microglia contacting the HRS-positive puncta, but it is unclear whether these puncta are secreted from neurons or engulfed by microglia before eventual internalization into microglia. To distinguish between these possibilities, we consulted the Allen Institute for Brain Science’s open-science 3D volumetric electron microscopy datasets. We first explored the 100 µm^3^ volume dataset from layer 2/3 of the visual cortex of a P36 male mouse^31^. The ssTEM dataset contains 364 segmented excitatory neurons and 25 segmented microglia, imaged at 3.58 x 3.58 x 40 nm resolution. The lengths of segmented microglial processes were manually surveyed. As reported previously, microglia engulfing partial or whole synapses that did not contain proteolytic organelles were observed (Extended Data Fig. 2), likely representing synaptic pruning. However, in numerous instances, proteolytic organelles in neuronal processes were engulfed by microglia (Fig. 4b, Extended Data Fig. 2, the x,y,z-coordinates of all instances are listed in Supplementary Table 1). Notably, microglia appeared to engulf a part of the presynaptic membrane containing proteolytic organelles, without disrupting the functional region of the synapse, leaving the active zone and post-synaptic density intact (Fig. 4a-b, c-d), as observed by superresolution imaging (Fig. 3). To distinguish between these two categories of microglial trogocytosis, we named this spatially specific enwrapping of proteolytic organelles, skoupocytosis, after the Greek word for garbage *skoupidia*.

**Fig. 4:**
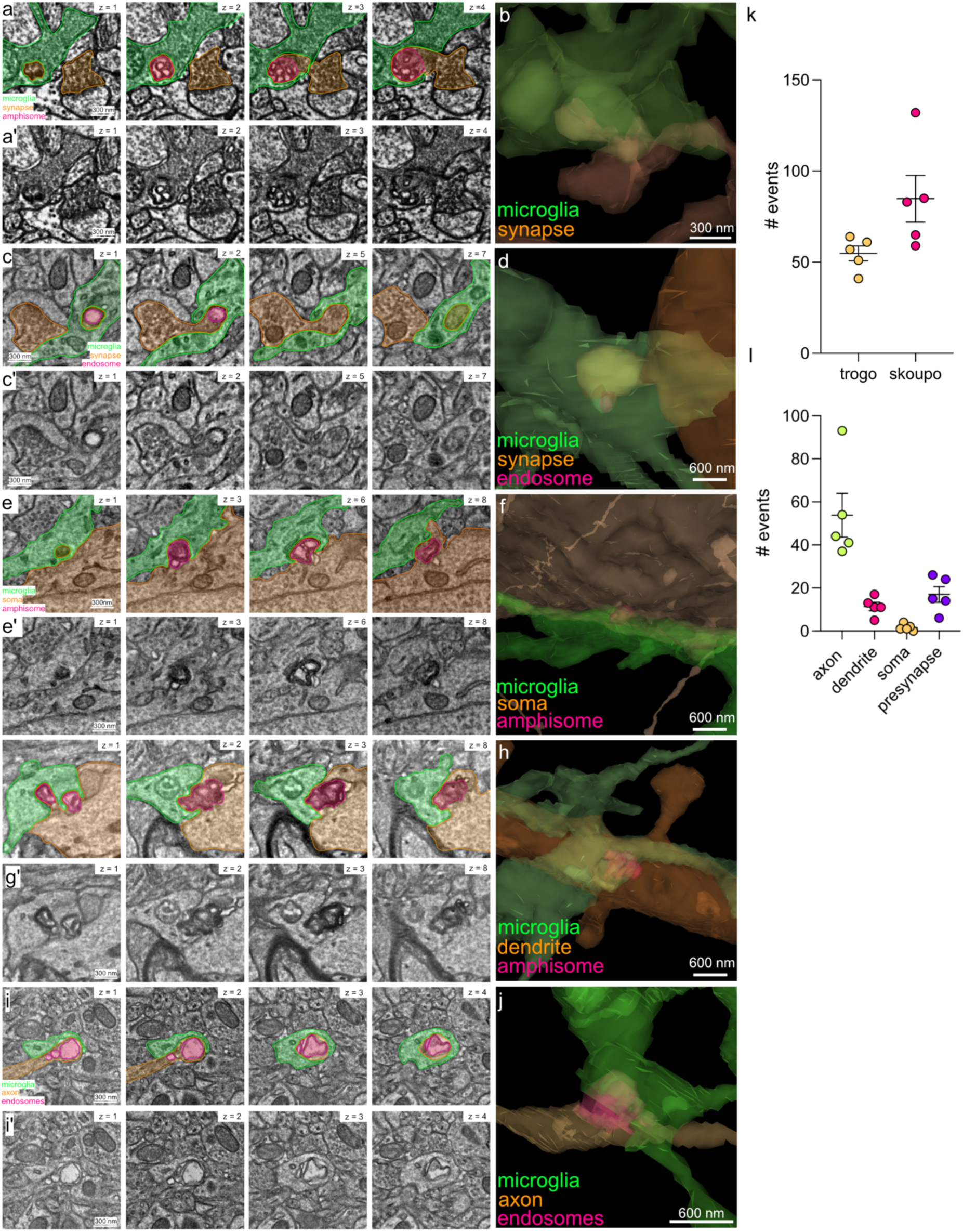
3D volumetric electron microscopy reveals spatially specific removal of proteolytic organelles from axons, dendrites, somas and synapses. (a) Representative slices of electron micrographs from P36 ‘pinky’ dataset showing microglial processes removing amphisome from presynapse. Microglia (green), amphisome (pink), neuron (orange). Scale bar 200 nm. (a’) same images without colored shades. (b) 3D rendered reconstruction of (a). Scale bar 600 nm. (c) Representative slices of electron micrographs from P87 ‘MICrONS’ dataset showing microglial processes removing large endosome from presynapse. Microglia (green), endosome (pink), neuron (orange). Scale bar 200 nm. (c’) same images without colored shades. (d) 3D rendered reconstruction of (c). Scale bar 600 nm. (e) Representative slices of electron micrographs from P87 ‘MICrONS’dataset showing microglial processes removing amphisome from soma. Microglia (green), amphisome (pink), neuron (orange). Scale bar 200 nm. (e’) same images without colored shades. (f) 3D rendered reconstruction of (e). Scale bar 600 nm. (g) Representative slices of electron micrographs from P87 ‘MICrONS’dataset showing microglial processes removing amphisome from dendrite. Microglia (green), amphisome (pink), neuron (orange). Scale bar 200 nm. (g’) same images without colored shades. (h) 3D rendered reconstruction of (g). Scale bar 600 nm. (i) Representative slices of electron micrographs from P87 ‘MICrONS’dataset showing microglial processes removing endosomes from axon. Microglia (green), endosomes (pink), neuron (orange). Scale bar 300 nm. (i’) same images without colored shades. (j) 3D rendered reconstruction of (i). Scale bar 600 nm. (k) Total number of microglial trogocytic events from 5 annotated microglia. Trogocytic events “trogo” (orange) or skoupocytic “skoupo” (pink). (l) Number of skoupocytic events per neuronal compartments: axons (green), dendrites (pink), soma (orange), and presynapses (purple).

To define the incidence of skoupocytosis, we categorized these interactions in the Allen Institute for Brain Science’s Machine Intelligence from Cortical Networks (MICrONs) dataset^32^. Taken from the visual cortex of a P87 mouse, the MICrONs dataset is 1.4 mm x .87 mm x .84 mm volume containing approximately 219,120 cells of which approximately 51,247 are neurons and a subset of 60 microglia have been identified and segmented. To get a snapshot of the types of interactions between microglia and neurons, 5 microglia from this dataset were surveyed and scored for the type of interaction and subcellular compartments. We classified trogocytic events as microglia enwrapping neurites where proteolytic organelles are not observed. Due to the static nature of this dataset, we only counted skoupocytic events where greater than 50% of the proteolytic organelle or membrane containing proteolytic organelle is enwrapped by microglial processes. Events where cell type, cell compartment or proteolytic organelle could not be identified due to incomplete segmentation, artefacts or inadequate resolution were excluded. An average of 55 putative trogocytic events and 85 putative skoupocytic events per microglia (Fig. 4i) were observed. On average 57% of putative skoupocytic events were enclosed by neuronal membrane while remaining events could not be determined due to limits in resolution (Extended Data Fig. 2c). Of note, putative skoupocytic events were found mostly in axons (54 events per microglia), followed by dendrites (11 events per microglia) (Fig. 4j). Similar events were also observed at the soma but at a much lower frequency (2 events per microglia). Of the incidences occurring at axons approximately a fifth of them occurred directly from presynapses (17 events per microglia), with only a small number of instances where some synaptic vesicles were included (Table S1). Together these data suggest skoupocytosis occurs ubiquitously across different cellular compartments of the neurons, but mostly in axons.

### ABHD16a accumulates near proteolytic organelles

The spatial specificity of skoupocytosis suggests that neurons may mark sites of proteolytic organelle accumulation to enable detection by microglia. Microglia have been shown to use immunological cues for phagocytosis and trogocytosis^26–28^. To identify potential cues, proteomic analysis was performed on isolated synaptic terminals (synaptosomes) (Fig. 5a). Previous studies have demonstrated that stimulating neurons pharmacologically with the GABA_A_ receptor inhibitor bicuculline and potassium channel inhibitor 4-aminopyridine (4-AP) increases the production of proteolytic organelles in presynapses, due to elevated usage of synaptic vesicles (Fig. 1)^16,33,34^. We reasoned that the presence of these proteolytic organelles may recruit molecular recognition components to the surface of presynapses for signaling to microglia. Cortical neurons at DIV 14 were treated with bicuculine and 4-AP for two hours. DMSO 0.1% treatment was used for control. Mass spectrometry was performed on the synaptosome fractions (Fig. 5a). After data were normalized across three independent cultures for quantitative analysis, 2549 unique proteins were identified (Fig. 5b, Supplementary Table 2). From these, 252 proteins had an abundance ratio greater than 1 and a p-value less than 0.05 (Fig. 5b, c), and 112 proteins are associated with the plasma membrane (Fig. 5d). One candidate protein, a/ß hydrolase domain-containing (ABHD16a), was of particular relevance, due to its interaction with phosphatidylserine (PS), a previously identified ‘eat me’ cue of microglia^35–37^. PS is a membrane phospholipid with two fatty acid tails and a serine headgroup residing on the inner leaflet of the plasma membrane; however, PS can be flipped to the outer leaflet by scramblase for phagocytosis by immune cells^28,38,39^. ABHD16a can cleave one of the fatty acid tails (sn-2) to generate lyso-PS^40–42^, which has been shown to induce debris clearance by microglia during demyelination^43^.

**Fig 5:**
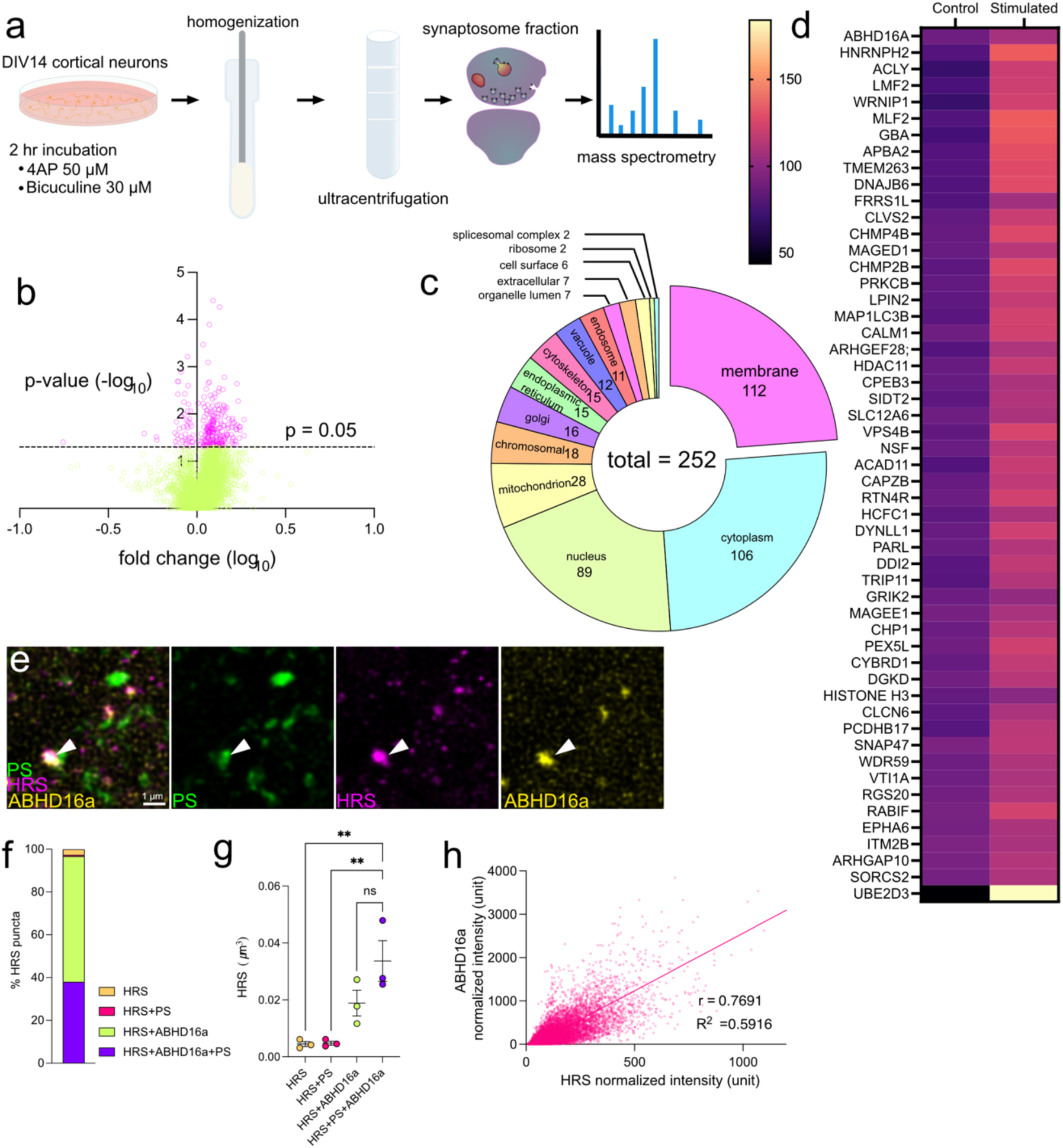
Phosphatidylserine lipase ABHD16a, and externalized PS colocalize with large proteolytic organelles. (a) Experimental design schematic: 1) DIV14 cortical neurons incubated with DMSO (control) or 50 µM 4AP + 30 µM bicuculline (treated), 2) neurons were homogenized, 3) ultracentrifugation, 4) synaptosome fraction isolated, 5) samples were analyzed using mass spectrometry. (b) Volcano plot of relative protein abundance between control and treated condition. 2549 unique proteins identified. Fold change (log10) versus the negative log10 of p-values. Significant changes are marked with a threshold (p = 0.05). (c) Pie chart of 252 identified proteins with >1 abundance ratio and p-value <0.05 differential expression between control and treated conditions based on cellular compartment. (d) Heatmap representation of 112 membrane proteins showing >1 abundance ratio p<0.05 differential expression between control and treated conditions. n=3 independent cultures. (e) Representative immunofluorescence images showing colocalization of HRS (dsRed to HRS-mCherry, magenta), ABHD16a (yellow), and externalized phosphatidylserine, PS (Annexin V, magenta) from acute brain slices prepared from P60 C57Bl6 mice n=3 mice. Scale bars represent 500 nm. (f) Percentage HRS puncta associated with PS and/or ABHD16a. (g) Comparison of volume (µm^3^) of HRS puncta, colocalized with PS and/or ABHD16a. Data shown as mean ± SEM. Statistical analysis ordinary one-way ANOVA with multiple comparisons (**p <0.001). (h) Scatter plot illustrating the correlation between normalized intensity values (arbitrary units, normalized to min/max) for HRS and ABHD16a, with a significant positive relationship indicated by the Pearson correlation coefficient (r² = 0.5916).

If lyso-PS is involved in skoupocytosis, ABHD16a and its substrate should accumulate near the HRS puncta. To test this possibility, acute slices were prepared from P60 *Cx3Cr1-eGFP* mice expressing neuronal HRS-mCherry, as previously described. Prior to fixation, slices were incubated with Annexin V-Alexa Fluor 647, which labels externalized PS (Fig.5e, f). SIM^2^ imaging revealed that nearly all HRS puncta colocalized with ABHD16a (96.0%). Approximately 40% of these HRS+ and ABHD16a+ puncta contain externalized PS. The rest of HRS puncta colocalized with PS (∼1%) or were isolated (3%). When HRS puncta contain ABHD16a, regardless of the presence of PS, the normalized (see Methods - Image Analysis) fluorescence intensity of HRS was higher compared to when they are not associated with ABHD16a (Fig. 5g) (Ordinary one-way ANOVA with Dunnett’s multiple comparisons, HRS vs HRS+PS+ABHD16a p = 0.0033, HRS+ PS vs HRS+PS+ABHD16a p = 0.0035). Further, there was a positive linear relationship between the intensity of HRS and ABHD16a (Fig. 5h) (Pearson Correlation r = 0.7691, R squared 0.5916). These data indicate that both the enzyme and its substrate are present near proteolytic organelles.

### ABHD16a is required for skoupocytosis

Previous studies have proposed lyso-PS can diffuse out from the membrane due to its amphipathic nature^44,45^. To test whether lyso-PS could act as a diffusible signal, unstimulated or stimulated (30 µM bicuculline, 50 µM 4AP, 1hr) DIV21 hippocampal neurons were treated with either 0.1% DMSO or KC01 (1 µM), a selective inhibitor for ABHD16a^41^, resulting in a significant decrease in lyso-PS production. Prior to fixation externalized PS was labelled with Annexin V-Alexa Fluor 594 (Fig. 6a-d). Annexin V only recognizes the head group and thus, cannot distinguish between PS and lyso-PS. Using superresolution SIM^2^ imaging, the ratio between externalized PS intensity and HRS puncta was calculated (Fig. 6e). In unstimulated conditions, the PS/HRS ratio was significantly increased in KC01-treated neurons (0.21) compared to DMSO-treated (0.09) (unpaired t-test with Welch’s correction, p = 0.0174, n = 4) (Fig. 6e). Following stimulation, there was also an increase in the PS/HRS ratio in KC01-treated neurons (0.42) compared to control (0.08) (unpaired t-test with Welch’s correction, p = 0.0583, n = 3) (Fig. 6e), suggesting that PS is retained in the outer leaflet of the plasma membrane when ABHD16a activity is blocked. This result implies that externalized PS would normally be processed to lyso-PS and leave the plasma membrane.

**Fig 6:**
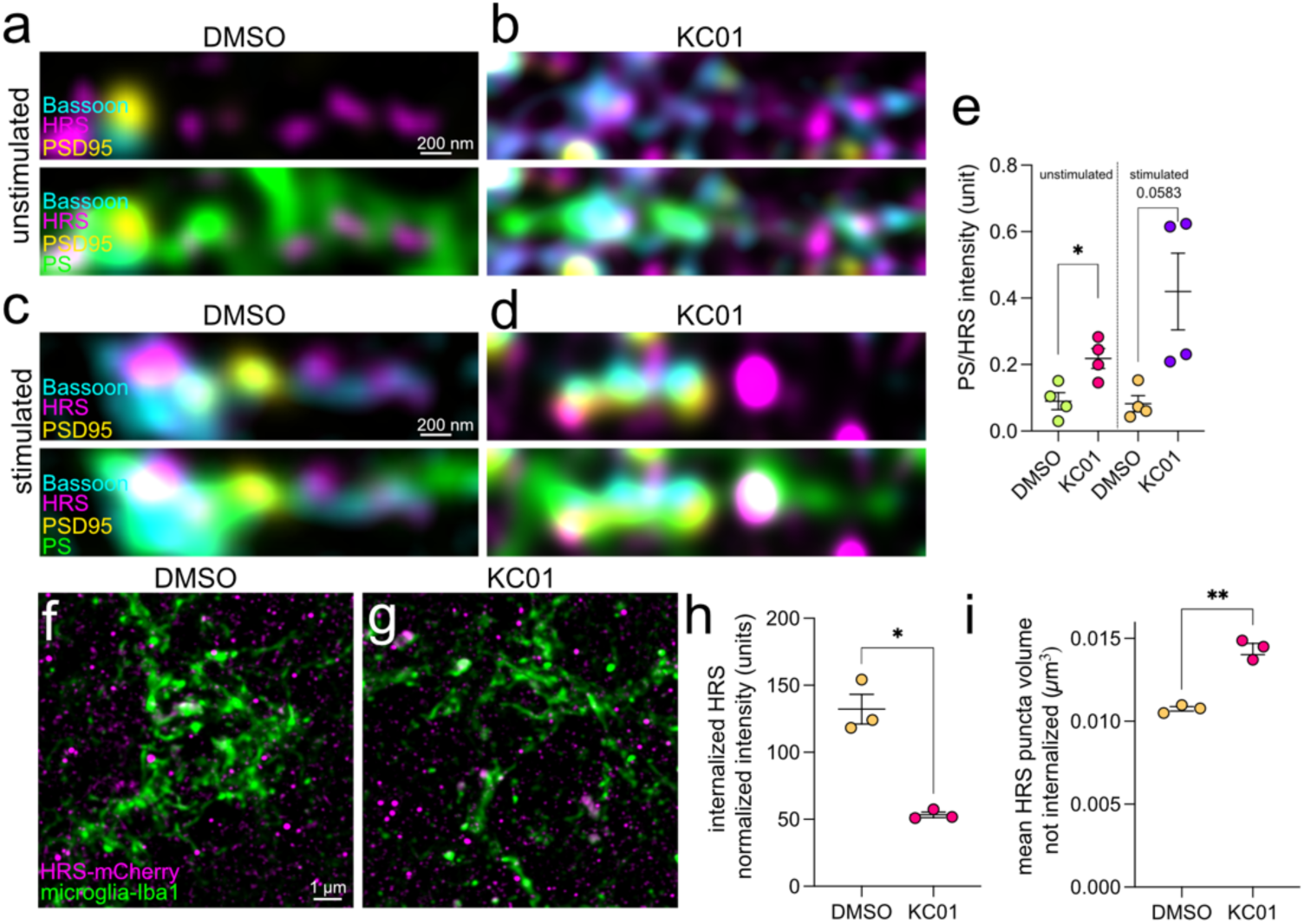
Phosphatidylserine (PS) lipase ABHD16a, and externalized PS co-localize with large proteolytic organelles. (a-b) Representative immunofluorescence images from DIV 21 hippocampal neurons treated with (a) DMSO (0.1%) or (b) ABHD16a inhibitor KC01 (1µM) for 2 hr. HRS antibody (magenta), PSD-95 (yellow), and presynaptic marker Bassoon (cyan) and externalized phosphatidylserine, PS (Annexin V, green). Scale bars represent 200 nm. (c-d) Representative immunofluorescence images from DIV 21 hippocampal neurons treated with (c) stim (50 µM 4AP, 30 µM bicuculline, DMSO 0.1%) or (d) stim + ABHD16a inhibitor KC01 (1µM) for 2 hr. HRS antibody (magenta), PSD-95 (yellow), and presynaptic marker Bassoon (cyan) and externalized phosphatidylserine, PS (Annexin V, green). Scale bars represent 200 nm. (e) Ratio of PS intensity to HRS puncta intensity in control and KC01 conditions. n = 4 independent cultures. Statistical analysis Welch’s t test (p = 0.0174). (f-g) Representative immunofluorescence images from acute brain slices neurons treated with (f) DMSO (0.1% control) or (g) ABHD16a inhibitor KC01 (1 µM) for 2 hr. Slices prepared from n=3 P60 C57Bl6 mice injected at P0 with PHP.eB.AAV.hSyn.HRS-mCherry. HRS (dsRed to HRS-mCherry, magenta), microglia-Iba1 (Iba1 antibody, green). Scale bars represent 1 µm. (h) Normalized HRS intensity internalized by microglia (arbitrary units, normalized to min/max) in control (DMSO 0.1%) or ABHD16a inhibitor KC01 (1µM). Data shown as mean ± SEM. Statistical analysis Welch’s t-test (* p < 0.05, ** p < 0.001). (i) Normalized HRS intensity outside of microglia (arbitrary units, normalized to min/max) in control (DMSO 0.1%) or ABHD16a inhibitor KC01 (1µM). Data shown as mean ± SEM. Statistical analysis Welch’s t-test (* p < 0.05, ** p < 0.001).

To determine whether ABHD16a activity is necessary for skoupocytosis, acute slices were treated with sham (0.1% DMSO) and 1 µM KC01, for 2 hours prior to fixation. HRS uptake by microglia was significantly reduced in KC01 treated slices (Fig. 6f) (unpaired t-test with Welch’s correction, p = 0.0169, n = 3) and correspondingly, the size of HRS puncta in neurons increased (Fig. 6g) (unpaired t-test with Welch’s correction, p = 0.0039, n = 3), suggesting that ABHD16a activity is necessary for skoupocytosis. Together, these results suggest that lyso-PS is used as a diffusible cue to initiate local proteolytic organelle removal by microglia through skoupocytosis.

## Discussion

Homeostatic microglia survey the entire brain parenchyma in under a few hours, contacting neurons and probing for changes to their microenvironment^46–49^. Microglia-synaptic contacts are well documented during development, where microglia remove silent synapses to fine tune circuits. This cellular pruning has also been implicated in learning and disease contexts throughout life^26–28,30^. This transfer of cellular constituents is typically inferred by the presence of synaptic proteins in microglia, suggesting large scale engulfment of synapses^50,51^. However, our findings demonstrate that microglia engage in the selective uptake of proteolytic organelles from neurites, including directly from presynapses, where proteolytic organelles are known to contain synaptic proteins^16,52^.

Whether neurons secrete proteolytic organelles or microglia actively engulf a part of the neuron that contain proteolytic organelles is not entirely conclusive. Microglia’s most basic function is as ‘garbage’ collectors for dead cells and debris. This function is crucial in disease and many studies have demonstrated the uptake of extracellular pathological aggregates such as ß-amyloid plaques, Tau and α-synuclein present in models of neurodegeneration^53–57^. However, how neurons deal with proteolytic organelles in the homeostatic mammalian CNS is less established. Neurons in the nematode *Caenorhabditis elegans* have been shown to secrete proteolytic organelles in the form of exophers, large membrane bound organelles containing protein aggregates and dysfunctional organelles, which are taken up by adjacent hypodermal cells and degraded^58,59^. Exophers are also produced by non-neuronal cells in mammals such as cardiomyocytes and proximal tubules in the kidneys^60,61^. However, exophers release by mammalian neurons has not been previously demonstrated. Recently, microglia have been shown to form tunneling nanotubes during neurodegeneration that facilitate the transfer of mitochondria and pathological protein aggregates such as α-synuclein^62–64^. However, while their presence *in vitro* has been extensively documented, the *in vivo* evidence is limited, and whether they occur in the non-diseased brain is unknown. While the possibility of these mechanisms occurring in skoupocytosis cannot be excluded; we favor a trogocytic form of interaction, where microglia recognize the parts of neurites that contain proteolytic organelles, due to the repeated examples that can be observed in the electron microscopy datasets. However, as static datasets it cannot be conclusively determined that these organelles are in fact internalized. Further studies to answer this question conclusively requires three color two photon microscopy, such as fluorescence lifetime imaging.

A remarkable trait of skoupocytosis is that microglia locate the areas of neurites containing proteolytic organelles and specifically extract them. To achieve this specificity, two major conditions must be met: 1. regions of accumulation must be able to communicate their internal state to nearby microglia through an external cue and 2. this cue must be spatially localized to restrict engulfment to proteolytic organelles. PS and lyso-PS signalling mediated by ABHD16a has the potential to fulfill both requirements. As proteolytic organelles form at synapses during synaptic activity, PS is known to accumulate on the surface of synapses through scramblase activity and has previously been suggested to provide localized cues for microglia during pruning^26–28^. Our data suggests that large HRS puncta are associated with both ABHD16a and externalized PS, and blocking of ABHD16a pharmacologically increases the amount of PS on the membrane. While some externalized PS is converted to lyso-PS, remaining PS could provide the localized cue for engulfment. However, at this stage it is unclear if PS is the ‘eat me’ signal for skoupocytosis, or whether the same downstream engulfment mechanisms like classic complement signalling or protective ‘don’t-eat-me’ signals are involved.

Lyso-PS is a conical lipid that promote curvature formation and can alter membrane mechanics^42^. Along this line, a recent paper shows that cell cortical tension regulates macrophage engulfment, where increasing cortical tension suppresses trogocytosis and promotes phagocytosis, suggesting softer membranes are required for trogocytosis^65^. However, the role this plays in skoupocytosis is yet to be explored.

Lyso-PS generated by ABHD16a from the outer leaflet may act as an attractant to guide microglia to the proteolytic burden synapses. Lyso-PS is a ligand for three known microglia GPCRs: GPR34, GPR174 and P2Y10R^44,45^. GPR34, the most abundant of these, is Gi-coupled, triggering PI3K-Akt and ERK signaling resulting in changes in microglia motility and promoting inflammatory cytokine release^43,66,67^. Lyso-PS sensed by GPR34 functions as chemoattractant in zebrafish brains by microglia during development, and lymphoid cells also use this pathway to find apoptotic neutrophils. Agonism of GPR34 promotes pathological aggregate clearance^53^, and myelin debris clearance in neurodegenerative models^43^. Further, GPR34 knock out mice have impaired phagocytosis^68^. However, GPR34 knockdown in the Alzheimer’s mouse model, APP/PS1, relieves cognitive deficits and suppresses inflammation^66^. More research is likely needed to disentangle how GPR34 signaling functions and contributes to neurodegeneration. The unexpected result of GPR34 knock out on neuroinflammation may be related to the purinergic G_12/13_-coupled receptor, P2Y10R in which activates RhoA and cytoskeletal remodeling, and suppresses inflammatory responses in microglia^69^. GPR174 is also G_12/13_-coupled in addition to G_s_-signaling but has mostly been detected in lymphoid tissues, specifically T cells^70^. Lyso-PS can also be incorporated into microglia inducing membrane remodeling and extension increasing their branching in culture independent of GPR34 and P2Y10R activity^71^. Lipid uptake by microglia would be advantageous to allow rapid membrane remodeling long distances from the soma with the added potential to also guide and promote membrane extension. Further work is needed to elucidate the downstream mechanisms.

The proteostatic requirements of neurons vary across cell type, brain region and age, and similar variety likely exists with skoupocytosis. Our data suggest differences between the somatosensory cortex and the hippocampus, potentially explained by differences in neural activity or neuron subtypes across regions. While organelle trafficking is difficult to observe *in vivo*, proteolytic organelles have been observed retrogradely trafficking in thalamocortical axons. It is hard to know however whether their final destination is degradation at the soma. Aly et al., 2025 observed the trafficking of amphisomes that were labelled in the distal thalamus reach the soma of these neurons suggesting there are likely some situations that require axonal trafficking^72^. Whether these requirements are cargo or neuron specific requires more exploration. The expression profiles of ABHD16a across brain regions and age support differences in proteostasis needs^73^. Proteomic studies enriching for synapses have demonstrated ABHD16a expression in the striatum, prefrontal cortex, hypothalamus, hippocampus and cerebellum^74^. Such widespread expression likely explains the impact dysfunction of ABHD lipases and disruption of lyso-PS processing has on CNS function. Mutations in ABHD12, responsible for metabolizing lyso-PS, presents as polyneuropathy, hearing loss, cerebellar ataxia, retinitis pigmentosa and cataracts (PHARC) a rare progressive neurodegenerative disease^40,41^.

Neuronal proteostasis and microglia are extricably linked in the pathophysiology of aging and neurodegeneration^35,75–78^. Declining neuronal proteolysis results in neuroinflamation which significantly impairs microglia function^75,77^. The degradative capacity of microglia decreases throughout life and a recent study shows that accumulation of synaptic proteins in microglia with aging can overwhelm microglia degraditive pathways^76^. In response, microglia release extracellular vesicles that propagate aggregate seeding, also engaging in abberrant synapse elimination accelerating cognitive decline^79,80^. Skoupocytosis provides a physiological pathway between neurons and microglia to help prevent deleterious protein accumulation during aging and neurodegeneration.

## Methods

### Animal husbandry and genotyping

All animal procedures were approved by the Institutional Animal Care and Use Committee (IACUC) under protocol number MO24M337 and adhered to the guidelines established by the National Institutes of Health. *C57BL/6N* and *Cx3Cr1-eGFP* mice were housed under standard conditions (22 ± 2°C temperature, 50 ± 10% humidity, 12-hour light/dark cycle) with ad libitum access to food and water. Mice were group-housed (up to 5 per cage) in individually ventilated cages (IVC) with environmental enrichment (e.g., nesting material and hut). Genomic DNA was extracted from ear clips using 50l lysis buffer (25 mM NaOH, 0.2 mM EDTA) at 100°C in heat block for 10 minutes. Mix was then neutralized with neutralization buffer (Tris-HCL 40 mM, pH 5.5) and 50 µl were used per reaction. PCR reactions were performed using a GoTaq G2 polymerase. Primer sequences can be found in supplementary methods. Amplified products were resolved on a 2% agarose gel containing ethidium bromide and visualized under UV light.

### Primary hippocampal and cortical neuron culture

Primary neuron culture was prepared as previously described^13^. Briefly, hippocampi or cortices were dissected from embryonic day 18 (E18) C57BL/6N mouse pups and transferred to ice-cold dissection media (Hanks Buffered Saline Solution, 1mM sodium pyruvate, 10mM HEPES, 30mM glucose, 1% penicillin-streptomycin). Under sterile conditions, meninges were carefully removed, and hippocampi were digested using 0.5 mg/ml papain and 0.01% DNaseI for 20 minutes at 37°C. Tissue was triturated using a fire-polished Pasteur pipette to obtain a single-cell suspension. Cells were plated at a density of 36,000 cells/cm² onto poly-L-lysine-coated coverslips (18 mm diameter, thickness #1.5) in Neurobasal medium supplemented with B-27 supplement, L-glutamine (2 mM), and penicillin/streptomycin (100 U/ml). Cultures were maintained in a humidified incubator at 37°C and 5% CO2. Half of the media was replaced every 3-4 days with glial-conditioned medium (supplementary materials). Experiments were performed at 14-22 days in vitro (DIV).

### Primary mixed glial culture preparation and harvesting

Mixed glial cultures protocol was adapted from Tamashiro, et al. 2012.^22^ Briefly, postnatal day 1-3 C57BL/6N or Cx3CR1-eGFP mice cortical tissues were dissected and mechanically dissociated in HBSS using a fire-polished Pasteur pipette. Cells were plated in T75 flasks (approximately 2 brains per flask) in Dulbecco’s Modified Eagle Medium supplemented with 10% FBS and penicillin/streptomycin (100 U/ml). Cultures were maintained at 37°C and 5% CO2 for 14 days. To obtain a purified microglial population, the bottom of T75 flask was vortexed gently for 2 minutes. The supernatant containing microglia was collected and centrifuged and cell pellet was resuspended in fresh DMEM and plated for further experiments.

### Lentivirus and adeno-associated virus production

Lentiviral particles were produced by transient transfection of HEK293T cells using the calcium phosphate precipitation method. The following plasmids were co-transfected pHR-CMV8.2 deltaR and pCMV-VSVG at a molar ratio of 4:3:2 and a plasmid containing the gene of interest (list of plasmids can be found in supplemental material). Briefly, 293T cells were plated at 70-80% confluency in T-75 flasks in Dulbecco’s Modified Medium supplemented with 10% FBS and penicillin/streptomycin. Supernatants containing lentivirus were collected at 72 hours post-transfection, filtered through a 0.45-μm filter, and concentrated using ultracentrifugation in Amicon Ultra 15 10K at 4°C. Viral titer was determined experimentally.

AAV particles were produced by transient transfection of HEK293T cells using protocol adapted from Negrini, et al 2020^81^. Capsid plasmid pUCmini-iCAP-PHP.eB and helper plasmid pAdDeltaF6 were co-transfected with plasmid containing gene of interest at 1:1:1 molar ratio. Briefly, 293T cells were plated at 70-80% confluency in T-75 flasks in DMEM. The DNA mix and containing PEI 1 μg/μl is added to cells. Supernatants containing AAV were collected at 24 hours and again at 72 hours post-transfection. Viral titer was determined using quantitative PCR using Ssoadvanced™ Universal SYBR with primers designed to ITR regions.

### Lentivirus and AAV infection of neurons and live cell imaging

Primary hippocampal neurons between DIV 6 and 14 were infected with AAV (serotype: PHP.eB; titer: 1 x 10^12^ vg/ml;). Virus was added directly to the culture media and were incubated with the virus at 37°C, 5% CO2. Half-media changes recommenced after 3 days of viral incubation. Cells used for live-cell imaging were transferred from their maintenance media into phenol red free Neurobasal media and incubated with JFX-646 for 30 minutes before washing with Neurobasal media once for 15 minutes. Coverslip was transferred to prewarmed imaging chamber (Warner Instruments, RC-49MFSH) and objective was heated (ALA Scientific Instruments, OBJ-HEATER). For trafficking imaging experiments Abbelight SaFe 180 was used with a 100x Nikon TIRF objective (NA = 1.49). Timelapse imaging at an interval of 200 ms, 50% laser power and 50% field of view illumination at using perfect focus to maintain focal plane. Post-processing using Abbelight software to correct for xy drift was used before image analysis (described below). For live imaging of microglia neuron co-culture Nikon Eclipse Ti2 wide-field microscope with 60x oil objective (NA =1.4) was used. Timelapse imaging was conducted at 1-minute intervals. Deconvolution was performed in ImageJ using PSF generator and Deconvolution lab 2^82–84^.

### Immunocytochemistry of cultured neurons

Neurons were fixed in pre-warmed 4% paraformaldehyde and 4% sucrose in PBS for 20 minutes at room temperature. Following fixation, neurons were washed three times with PBS and permeabilized with 0.1% Triton X-100 in PBS for 10 minutes. Non-specific binding was blocked with 1% BSA in PBS for 30 minutes at room temperature. Neurons were incubated with primary antibodies diluted in blocking solution overnight at 4°C. After washing three times with PBS, neurons were incubated with secondary antibodies (list of primary and secondary antibodies and their concentrations can be found in supplemental materials). Coverslips were mounted onto glass slides using ProLong Diamond antifade mountant and cured for 48 hours at room temperature. Cells were imaged using Zeiss Elyra 5 and SIM^2^ post processing was applied before image analysis as described below.

### P0 injections

P0-2 from Cx3Cr1-eGFP pups were cryo-anaesthetized on ice and injection site is sterilized using 70% ethanol prior to injections. Mice were injected with 2 µl of PHP.eB (titer: 1 x 10^12^ vg/ml) using a Hamilton syringe at coordinates: AP: -0.5 mm, ML: ±1.0 mm, DV: -2.5 mm from bregma. Mice were then returned to heat pad for recovery before returning to home cage.

### Cardiac perfusion and immunohistochemistry of brain tissue

Mice were deeply anesthetized with Avertin (200-500 mg/kg) (protocol in supplementary materials) and administered intraperitoneally. Once the absence of a toe-pinch reflex was confirmed, mice were transcardially perfused with 4% PFA. Brains were carefully removed followed by 2-hour post-fix in 4% PFA in PBS. Brains were transferred to 30% sucrose in PBS overnight at 4°C. Tissue was then transferred to cryo-molds and OCT added before freezing in a weight boat suspended on liquid nitrogen and stored at -80°C.

Brains were sectioned coronally at 30 µm thickness using a cryostat. Free-floating sections were incubated with blocking solution (1% BSA, 0.2% Triton X-100 in PBS) for 1-hour on rotating platform. Sections were incubated with primary antibodies diluted in blocking solution for 24 hours at 4°C with gentle agitation on a rotating platform. After 3x PBS washes, sections were incubated with secondary antibodies for 24 hours at 4°C. Sections were then mounted onto glass slides using ProLong Diamond antifade and cured for 48 hours at room temperature. Imaging of fixed brain slices was completed using Zeiss Elyra 5 or SoRa Nikon.

### Correlative electron microscopy

Neurons were cultured on sapphire disks (#616-100; Technotrade) that were carbon-coated in a grid pattern (using finder grid masks, #16770162; Leica microsystems). After 1 mg/ml poly-Lysine (#P2636; Sigma-Aldrich) coating overnight, neurons were seeded at a density of 2.5 × 10^2^ cell/mm^2^ in 12-well tissue-culture treated plates (#3512; Corning) and cultured in NM0 medium (Neurobasal medium (#21103049; Gibco)) supplemented with 2% B27 plus (# A3582801; Life Technologies) and 1% GlutaMAX (#35050061; Thermo Fisher Scientific). HRS-Halo and vGlut1-pHluorin were transduced by lentivirus at DIV 6, and experiments performed at 14–15 DIV. To induce neuronal activity, neurons were incubated with 40 μM bicuculline (Bic, #0130/50; Tocris) and 50 μM 4-aminopyridine (4AP, #940; Tocris) at 37°C. After 2 hours of incubation, half of the medium was changed to NM0 with 200 nM Jenelia-Halo-549 nm (#SB-Jenelia-Halo-549; Lavis Lab) and incubated for an additional 30 min at 37°C. Neurons on the sapphire disks were washed 3 times with PBS, fixed with 4% paraformaldehyde (#15714; EMS) and 4% sucrose (#S0389; Sigma-Aldrich) in PBS for 20 min at room temperature, then washed 3 times with PBS.

For the confocal imaging, sapphire disks where flipped upside down on an 18-mm coverslips mounted on a chamber (#MS-508S; ALA Science) filled with PBS. Z-series of fluorescence images were acquired using Zeiss LSM 880 confocal microscope equipped with a 40× objective lens at 2,048 × 2,048 pixel resolution. Bright field images of the carbon grid pattern and neurons on the sapphire disks were also acquired. Following imaging, neurons were post-fixed with fixation buffer 1 (2% glutaraldehyde, 1 mM CaCl_2_, 0.1 M Sodium cacodylate, pH 7.4 in dH_2_O) for 1 hour on ice. Samples were washed 3 times using 0.1 M Sodium cacodylate then fixed in fixation buffer 2 (1% osmium tetroxide, 1% potassium ferrocyanide in dH_2_O) for 1 hour on ice. Cells were then washed 3 times with dH_2_0 before staining with 2% uranyl acetate in water for 30 minutes at room temperature. To dehydrate samples, they were incubated in a graded series of ethanol (50%, 70%, 90%, 100%) for 5 minutes each at room temperature. Final 100% ethanol incubation was done 3 times before embedding in 100% epoxy resin at 4°C overnight. Samples were then transferred to new 100% epon 3 times for 2 hours each before curing in 60°C oven for 48 hours. Approximately 40 ultrathin serial sections (80 nm) were cut using an ultramicrotome (UC7; Leica) and mounted on to piliform-coated copper grids. Sections were imaged using a Hitachi 7600 transmission electron microscope. The confocal and electron microscopy images were roughly aligned based on size of the pixels and magnification, and the carbon-coated grid patterns. The alignment was slightly adjusted based on the visible morphological features.

### Cranial window implantation and *in vivo* 2-photon imaging

Adult (8-12 weeks old) Cx3CR1-eGFP mice were deeply anesthetized with isoflurane 5% in oxygen and maintained at 1-2% isoflurane in oxygen. Mice were given an injection of lidocaine 0.5% (7 mg/kg) injection under the scalp, dexamethasone injection (1 µg/g, 0.001 mg/kg), and 1 ml of sterile saline injection prior to surgery and maintained on a heat pad at 37°C during the surgery. Hair was shaved and hair removal cream used before sterilizing scalp using 70% ethanol, iodine and 70% ethanol again. A craniotomy minimally larger then 3 mm in diameter was performed over the somatosensory cortex (coordinates: Bregma +0.5 mm, lateral 3 mm) using a dental drill and periodically cooled using chilled saline. A coverslip (3 mm diameter, #1;) was cemented in place using dental cement (C and B Metabond). A custom-made headplate was attached to the skull using dental cement to allow for head fixation during imaging. Mice were given carprofen (10 mg/kg) immediately and daily for 3 days after surgery for pain relief.

For imaging, mice were anesthetized with 5% isoflurane in oxygen and transferred to imaging stage. Anesthesia was maintained using 1-2% isoflurane in oxygen. Custom headplate was used to head-fix mouse for imaging, and body temperature was maintained using heat pad for the session. *In vivo* two-photon imaging was performed using Thorlabs Bergamo II microscope equipped with a Discover NX laser and a 25x water-immersion objective (Nikon, model: XLPLN25XWMP2). Imaging was performed at 920 nm wavelength using galvo-resonance scanning at a depth of 100-150 μm from the surface. Deconvolution was performed in ImageJ using PSF generator and Deconvolution lab 2^82–84^. Motion correction using Correct 3D Drift plugin in ImageJ was used to register between channels^85^.

### Construct cloning

List of constructs cloned for this project can be found in supplementary material. One of two methods was used to generate new constructs; Takara bio infusion or restriction enzyme digestion and ligation (NEB, relevant enzymes and buffers used as per manufacturer’s instructions). Successful cloning was confirmed by whole plasmid sequencing.

### Synaptosome preparation, mass spectrometry and proteomic analysis

Cortical neurons were seeding at a density of 8 million cell/plate on PLL coated 10 cm^2^ plates. At DIV14 cells were treated with either dimethyl sulfoxide (DMSO, 0.1%) or 4-aminopyridine (4-AP, 50 μM) and Bicuculine (30 μM) for 2 hours. After incubation cells were scraped using a cell scraper and transferred to chilled 7 ml Dounce homogenizer and homogenized in 6 volumes of ice-cold homogenization buffer (0.32 M sucrose, 5 mM HEPES, 1x cOmplete protease inhibitor, pH 7.4) until the suspension appeared uniform. An aliquot (10 μl) of the homogenate was diluted 1:10 in homogenization buffer and retained for sample analysis. The homogenate was transferred to a pre-cooled tube and centrifuged at 1,000× g for 10 min at 4°C (SS-34 rotor). The supernatant (S1) was moved to a fresh tube and a 1:10 dilution (10 μl + 90 μl) was retained on ice for sample analysis. The pellet (P1) was estimated by comparison to a reference volume, resuspended in 20× its pellet volume in homogenization buffer, and an aliquot (100 μl) was retained on ice for sample analysis. A discontinuous Ficoll (13%, 9%, 6%) gradient was prepared in an SW-41Ti ultracentrifuge tube by layering (bottom to top) 1 ml 13% Ficoll, 1 ml 9% Ficoll, and 1 ml 6% (all in 0.32 M sucrose 5 mM HEPES, 1x cOmplete protease inhibitor, pH 7.4). The S1 suspension was layered on top of the gradient, taking care not to disturb the interfaces. Gradients were centrifuged at 82,500× g for 35min at 4 °C in an SW-41Ti rotor. After centrifugation, a thin, hazy band (synaptosomes) located at the interface between 19% and 13% Ficoll was collected (∼30 µL) by gentle pipetting. The collected synaptosome fraction was diluted with 3x volumes of 0.32 M sucrose 5 mM HEPES, 1x cOmplete protease inhibitor (pH 7.4) and transferred to a 1 ml Eppendorf tube. Synaptosomes were pelleted by centrifugation at 6,000× g for 3 min at 4°C. The supernatant was removed, and the pellet was resuspended in 100 μl 80 mM Tris-HCl, pH 8.0. Resuspended synaptosomes were stored at −80°C before being sent for mass spectrometry. Briefly, the mass spectrometry consisted of Multi-Dimensional Protein Identification Technology (MuDPIT) basic C18 reverse phase (bRP) chromatograph followed by acidic C18 reverse chromatography with Tandem Mass Tags (TMT, Thermo Scientific 90309, lot#TK255306) for chemical labeling with isobaric mass tags. Proteomic analysis was conducted using Proteome Discovery version 2.4 and Mascot 6.2.6 and the RefSeq2021_204_mus_musculus protein database used to identify peptides. Mass tolerance for precursor ions was set to 5 ppm, and mass tolerance for product ions set to 0.02 Da.

### Acute slice preparation

Mice were deeply anesthetized with Avertin (125-150 mg/kg) administered intraperitoneally. Once the absence of a toe-pinch reflex was confirmed, mice were transcardially perfused with sucrose cutting solution (195 mM Sucrose, 10 mM NaCl, 2.5 mM KCl, 1.25 mM NaH_2_PO_4_, 26.2 mM NaHCO_3,_ 15 mM Glucose). Brains were quickly in ice-cold sucrose cutting solution and mounted for vibratome sectioning using super glue. Coronal slices (1 mm) were prepared using a vibratome or tissue chopper. Slices were incubated in a recovery chamber containing artificial cerebral spinal fluid (ACSF, 124 mM NaCl, 2.5 mM KCl, 1.4 mM NaH_2_PO_4_, 30 mM NaHCO_3_, 25 mM HEPES, 25 mM Glucose) at 32°C for 30 minutes before incubation with KC01 at 1 μM diluted from 1mM stock solution dissolved in DMSO. Slices were fixed with 4% PFA for 2 hours at room temperature after drug treatment and incubated with sucrose 30% overnight before embedding to be cryosectioned as described above.

### Image analysis

Image analysis was performed using Fiji/ImageJ (v. 2.3.0) and Icy using a combination of open-source plugins and custom-written macros and code. For analysis of organelle tracking time-lapse imaging, ROIs were generated using Wavelet Spot Generator plugin and tracking was done using the Spot Tracking Plugin^18,86,87^. This method generates a multiscale dataset by decomposing the original image using *à trous* wavelet transform, thresholding out non-significant coefficients^86^. By correlating information from different levels of resolution, this allows enhanced peak detection and noise reduction^86^. For analysis of microglia internalization *in vitro*, ROIs were manually segmented using differential interference contrast, and intensity measurements were taken using ImageJ and normalized using min/max normalization. For analysis of microglia internalization *in vivo*, ROIs were generated using Wavelet Spot Generator plugin. From these segmented images, stationary organelles were manually selected, and 3D ROI volume was extracted across the video. Intensity measurements were normalized using min/max normalization. To filter ROIs for overlap either to exclude ROIs that were internalized in microglia or partially intersecting microglia (Figs. 2,4,6), logical operation statements using Icy protocols. ROIs considered internalized by microglia must contain 100% overlap of pixels with microglia signal, while intersecting ROIs are required to have greater than 1 pixel overlap with microglial ROIs.

### Volumetric electron microscopy analysis

Dataset is open-access available at https://www.microns-explorer.org/ from the Allen Institute for Brain Science’s Machine Intelligence from Cortical Networks (MICrONs) P87 dataset^32^. Processes of 9 microglia were manually surveyed and scored for the type of interaction and cellular compartment. We classified trogocytic events as microglia processes enwrapping neurites where proteolytic organelles were not observed. Skoupocytic events where greater than 50% of the proteolytic organelle or membrane containing proteolytic organelle was internalized in microglia. Events where cell type, cell compartment or proteolytic organelle could not be identified due to incomplete segmentation, artefacts or inadequate resolution were excluded.

### Statistical analysis

All culture experiments have 3 independent biological replicates from cultures done on different days. Tests for normality were conducted on the data prior to testing for statistical significance to determine which type of test was performed. For live imaging of organelle trafficking, 1 cell from 3 coverslips per independent culture days were averaged and statistical analysis was compared using independent replicates. For microglia fixed *in vitro* imaging analysis, 6-16 microglia were imaged randomly from 3 independent biological replicates from cultures done on different days. For *in vivo* imaging, n = 6 mice were imaged at an interval of 1 minute, and manually analyzed for instances of skoupocytosis (see Data Availability). For fixed imaging, 7-20 microglia approximately 20-30 µm in z-depth, were imaged randomly from across the somatosensory cortex and the hippocampus, the mean of these images of n = 3 mice were compared for statistics. For analysis of HRS density and presence at synapses, 3 images with a z-depth of x µm were analyzed and the mean of n = 3 mice compared. For proteomics, Proteome Discovery version 2.4 was used to visualize data. The FDR was set to 1%. Proteins considered enriched had a p-value greater than 0.05, and a Log_2_ fold change greater than 0.26, and proteins considered depleted had a p-value greater than 0.05, and a Log_2_ fold change less than -0.26. All analysis was completed using GraphPad Prism 10.

## Acknowledgements

We would like to thank Clarissa Waites for gifting HRS-mCherry constructs. We also thank Barbara Smith, Mike Delanoy, LaToya Roker, Loza Lee, and Scot Kuo at the Johns Hopkins Microscopy Facility for technical and administrative assistance in confocal microscopy; Aleksandr Smirnov and Susan McTeer at the Johns Hopkins Neuroscience Imaging Center for technical and administrative assistance in 2-photon microscopy; Robert Cole at the Johns Hopkins Biological Chemistry Mass Spectrometry and Proteomics Core.

## Author contributions

R.P., J.G. and S.W. conceived the study and designed the experiments. S.W. oversaw the overall project and funded the research. R.P. performed live-cell organelle tracking experiments and analysis. R.P. and J.G. performed live-cell, R.P., J.G., and E.P fixed cell *in vitro* microglia experiments. Y.I. performed correlative electron microscopy experiments and analysis. R.P., F.M.D, S.S, and J.B. performed volumetric EM analysis. R.P. performed cranial window surgeries and *in vivo* 2-photon imaging. Y.X. and D.B. provided technical assistance for 2-photon imaging. J.G. performed proteomic experiments and analysis. R.P., F.M.D, E.P. and L.T. performed intracerebroventricular injections, acute-slicing experiments and imaging. R.P. and S.W. wrote the manuscript. All authors edited the manuscript.

## Materials and correspondence

Professor Shigeki Watanabe |, Johns Hopkins School of Medicine, Department of Cell Biology, 725 N Wolfe St, Baltimore, MD, USA, 21205

---

**Extended Data Fig 1:**
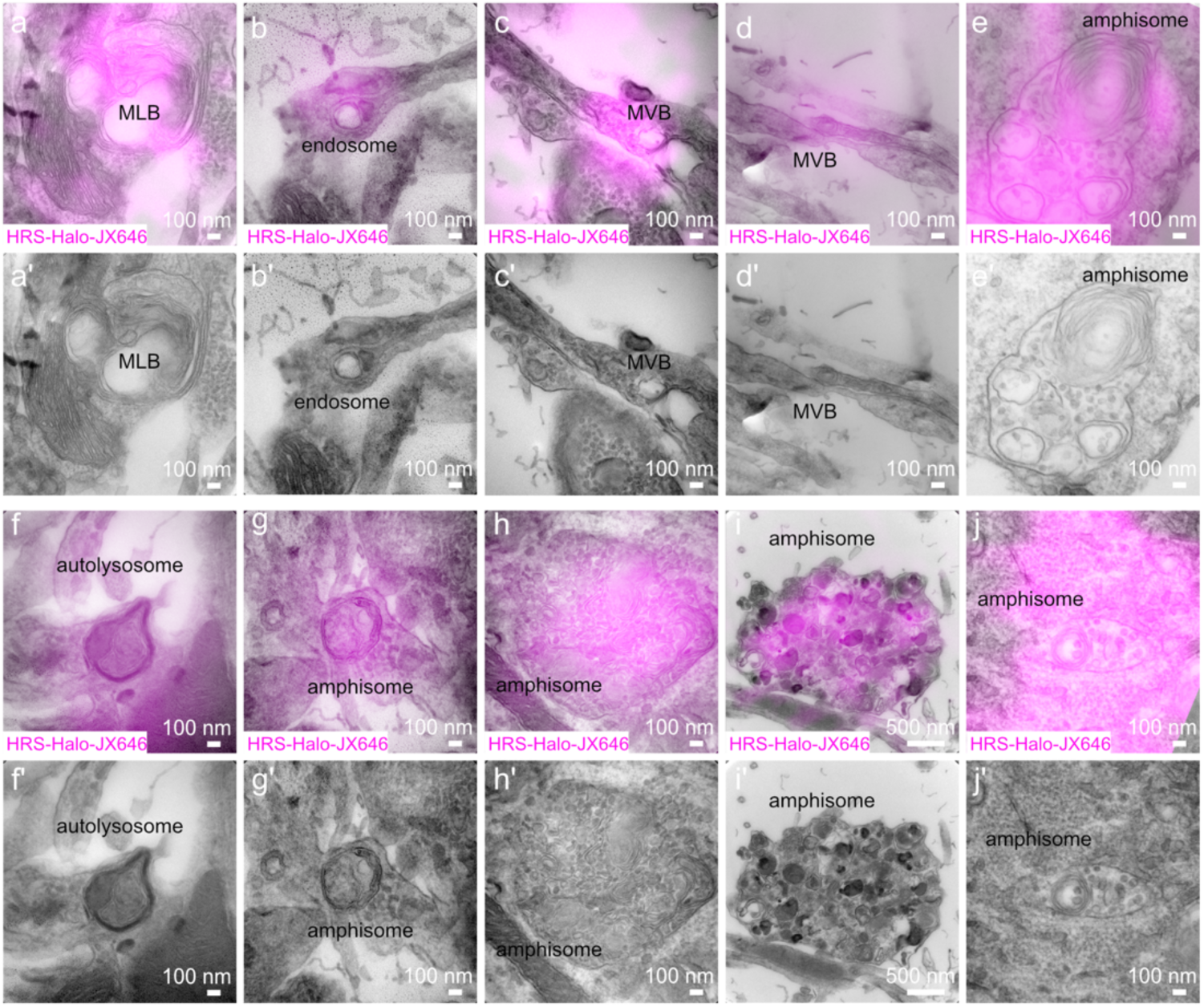
(a-j) Electron micrographs overlayed with correlative fluorescent signal from HRS-HaloTag-JFX646-labeled organelles: (a) multi-lamellar body, (b)endosome, (c) multi-vesicular body, (d-e) amphisome, (f) autolysosome, (g-j) amphisome. Scale bars are 100 µm for (a–h, j) and 500 µm for (i). n = 3 independent hippocampal cultures.

**Extended Data Fig. 2:**
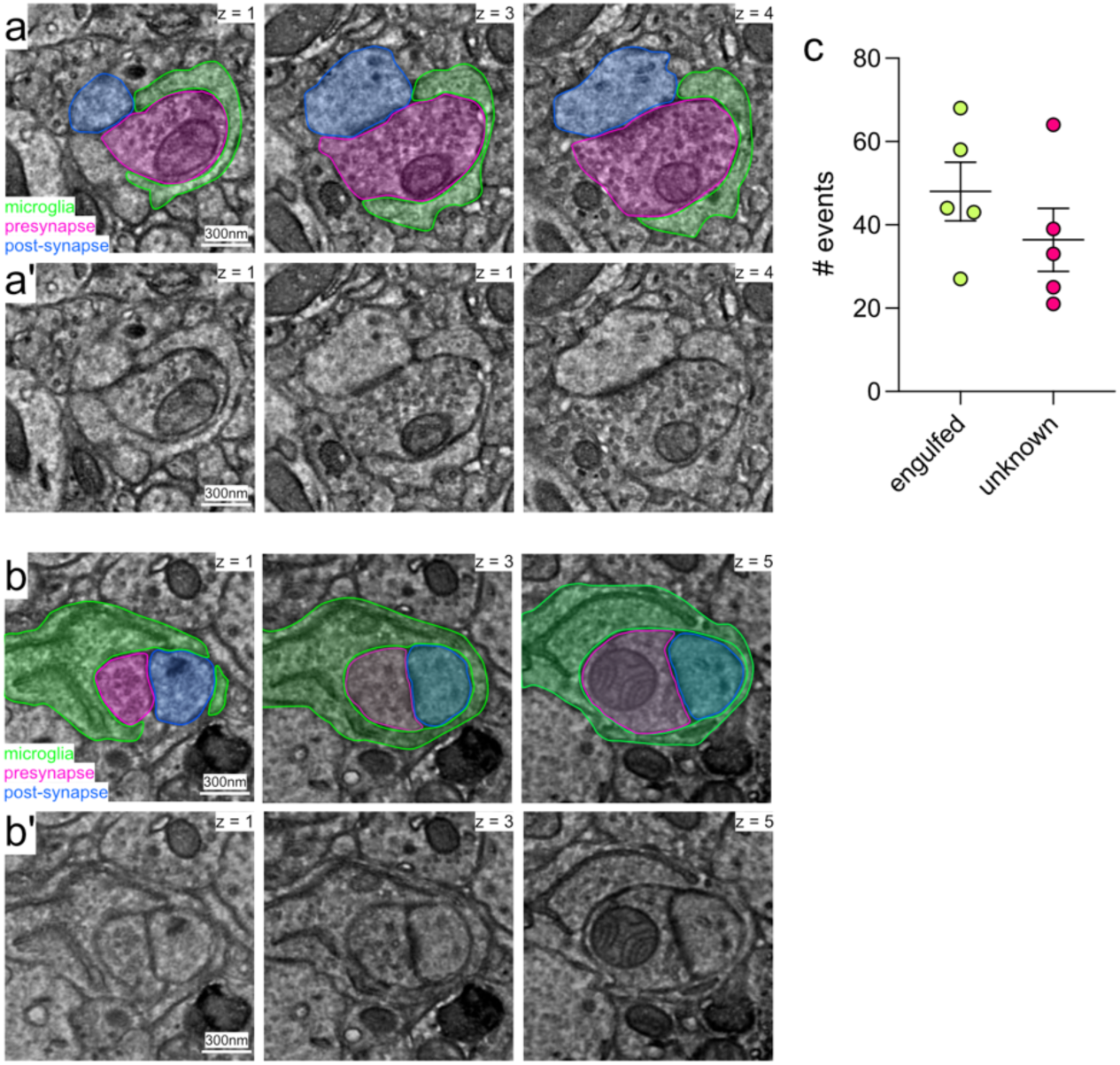
(a) Representative slices of electron micrographs from P87 ‘MICrONS’ dataset showing microglial processes removing presynapse. Microglia (green), presynapse (magenta), post-synapse (blue). Scale bar 300 nm. (b) Representative slices of electron micrographs from P87 ‘MICrONS’ dataset showing microglial processes removing whole synapse. Microglia (green), presynapse (magenta), post-synapse (blue). Scale bar 300 nm. (c) Number of microglial skoupocytic events where 1) engulfment of proteolytic organelles with intact neuronal membrane or 2) unknown whether neuronal membrane is intact. Taken from 5 annotated microglia. Engulfed events “engulfed” (green) or unknown “unknown” (pink).

<u>Supplementary Video 1: Microglia internalize neuronal proteolytic organelles in vitro</u>

(a) Time-lapse video of microglia from Cx3Cr1-eGFP (green) taking up neuronal HRS-mCherry^+^ (magenta) organelles. 1-minute imaging intervals. Scale bars represent 1 µm. Representative images from P70 mouse. Images selected from n = 8 mice.

<u>Supplementary Video 2a: Proteolytic organelles from neurons internalized in microglia *in vivo*</u>

(a) Time-lapse video of microglia from Cx3Cr1-eGFP (green) with HRS-mCherry^+^ (magenta) organelles internalized in microglia. 1-minute imaging intervals. Scale bars represent 1 µm. Representative images from p70 mouse.

<u>Supplementary Video 2b: Large proteolytic organelles are stationary in neurons *in vivo*</u>

(b) Time-lapse video of microglia from Cx3Cr1-eGFP (green) taking up neuronal HRS-mCherry^+^ (magenta) organelles. 1-minute imaging intervals. Scale bars represent 1 µm. Representative images from P70 mouse.

<u>Supplementary Video 2c: Microglia take up proteolytic organelles from neurons *in vivo*</u>

(c) Time-lapse video of microglia from Cx3Cr1-eGFP (green) taking up neuronal HRS-mCherry^+^ (magenta) organelles. 1-minute imaging intervals. Scale bars represent 1 µm. Representative images from P70 mouse.

<u>Supplementary Video 2d: Microglia take up proteolytic organelles from neurons *in vivo*</u>

(d) Time-lapse video of interactions between microglia from Cx3Cr1-eGFP (green) mouse with neuronal HRS-mCherry (magenta) labeled organelles. 1-minute imaging intervals. Scale bars represent 500 nm. Representative images from P70 mouse.

<u>Supplementary Video 2e: Microglia take up proteolytic organelles from neurons *in vivo*</u>

(d) Time-lapse video of interactions between microglia from Cx3Cr1-eGFP (green) mouse with neuronal HRS-mCherry (magenta) labeled organelles. 1-minute imaging intervals. Scale bars represent 1 µm. Representative images from P75 mouse.

